# Deep Learning-enabled Sperm Morphology Analysis of Bovine Sperm for label-free Imaging Flow Cytometry

**DOI:** 10.1101/2025.06.12.659335

**Authors:** Anel Umirbaeva, Andrey Kurenkov, Bolat Seisenov, Kanat Zhumanov, Kadyrbay Tajiyev, Mirzhan Mustafin, Ivan A. Vorobjev, Natasha S. Barteneva

## Abstract

Data analysis of sperm morphology is critical for evaluating bull fertility, yet it is often performed using light microscopy and staining techniques in a highly subjective and manual manner. In this study, we introduce a scalable, high-resolution approach combining label-free Imaging Flow Cytometry (IFC) with deep learning for automated classification of bovine sperm morphology. We analyzed 436,374 single-cell images obtained from three prominent bull breeds in Kazakhstan - Kazakh Whitehead, Auliekol, and Simmental from fresh and cryopreserved sperm - providing a uniquely large and diverse dataset. The dataset was used for training and evaluation of deep learning models, among which the convolutional neural network (CNN) MobileNetV4 yielded superior results, achieving 92.3% accuracy and a 0.91 F1-score after training with a Layer-wise Pretraining and Fine-Tuning (LP-FT) strategy. The model classified spermatozoa into eight distinct morphological categories. The CNN-based pipeline ensured consistent, observer-independent classification across all samples. Testing across different conditions and breeds resulted in a 5-10% drop in generalization performance, highlighting the impact of domain-specific biases and underscoring the need for larger, standardized datasets. The proportion of morphologically abnormal spermatozoa varied between seasons and after cryopreservation. This study highlights the advantages of integrating IFC and artificial intelligence (AI) algorithms for robust, high-throughput, and objective label-free spermatozoa morphology assessment in fresh and cryopreserved sperm, offering a promising tool for improving fertility diagnostics and breeding strategies in veterinary practice.

## 1. Introduction

Successful oocyte fertilization relies on high sperm quality, which is reflected in sperm cell count, motility, morphological structure, and functionality (1–4). Accurate, multi-parametric evaluation of these quality indicators is critical for the efficiency of assisted-reproduction technologies (5–6). The routine evaluation of semen has traditionally included the assessment of morphological abnormalities (7–9); however, it is challenging and primarily relies on staining techniques (10–11). Sperm cryopreservation and semen refrigeration have been employed for preserving sperm fertilizing function, and methods for assessing and predicting sperm function are critical for the optimization of fertility outcome (12).

However, even with the seemingly widespread success of cryopreservation and sperm banking, it is not completely understood how to improve spermatozoa cryosurvival and decrease damage. The increased percentage of sperm abnormalities after freezing includes coiled and bent spermatozoa tails (13), cracked tails, and detached heads (14). The most pronounced damage happens in the sperm plasma membrane after the freeze-thawing step.

Morphological abnormalities are often correlated with other biomarkers of sperm function (15–17). For example, head defects have been linked to the altered expression of proteins involved in sperm capacitation and sperm-egg interaction (18) and chromatin packaging anomalies that may compromise nuclear membrane integrity (19). Tail abnormalities are commonly associated with impaired motility and mitochondrial dysfunction (20–21), and severe flagellar deformabilities can result in loss of motility and infertility. Other morphological abnormalities, such as the presence of excess residual cytoplasm, may indicate incomplete maturation of cells or high osmotic stress during storage or transportation, ultimately reducing viability (22). Imaging flow cytometry (IFC) is one of the few technologies capable of simultaneously analyzing multiple functional parameters coupled with morphological assessment at the single-cell level (23). Considering the complex structure of spermatozoa and the frequent occurrence of multiple co-existing abnormalities within a single cell, traditional manual morphological assessment, which is a long-standing gold standard for identifying structural anomalies (24), has significant limitations in accurate categorization of images.

Biomarkers discovery efforts in animal and human sperm selection are also closely related to the adaptation of conventional and imaging flow cytometry (FC and IFC) (16, 25–30). Artificial intelligence (AI) algorithms, in combination with advanced imaging technologies, could potentially provide the solution to the efficiency and subjectivity challenge of sperm quality assessment and selection (31–34). IFC is, thus, a suitable option for high-throughput sperm analysis, resulting in tens of thousands of label-free single-cell events with about a thousand morphological parameters. Such large image datasets allow the training of the AI models for faster and more precise sorting into morphological categories (35).

The primary goal of this research was to develop a high-performance deep-learning model for classifying bovine sperm morphology, both before and after cryopreservation. This was done by creating a large IFC dataset comprised of fresh and frozen semen samples from three local bull breeds: Kazakh Whitehead, Simmental, and Auliekol. Additionally, the study aimed to go beyond standard model accuracy metrics by conducting systematic cross-validation experiments for the evaluation of the model’s robustness and generalizability across varying sample preparation conditions and cattle breeds. The findings from these analyses provide valuable insights into the potential application of this model in biological and veterinary contexts.

## 2. Materials and Methods

### 2.1. Animals and semen collection

Sperm from six beef bull sires (4 Kazakh Whitehead, 1 Simmental, and 1 Auliekol) were used in this study. Cryopreserved straws of 6 qualified ejaculates (motility > 60%) of bulls obtained during the spring-summer season and diluted in OptiXcell^®^ (IMV Technologies, France) and fresh ejaculates from these bulls were used for this study. The sperm were not collected specifically for research but were a byproduct of the standard production process and, therefore, exempt from Nazarbayev University Institutional Animal Care and Use Committee (IACUC) oversight. All procedures in the “Asyl-Tulik” facility were performed in accordance with relevant animal procedural guidelines to ensure animal welfare and conformance to ethical integrity regulations.

### 2.2. CASA analysis of motility

Computer-assisted sperm analysis, or CASA, was performed by the Hamilton Thorne Motility Analyzer HTM-2030 (Hamilton Thorne LTD., USA) to evaluate sperm motility values, including cell count, progressive motility (PM, %), total motility (TM, %), elongation (μm), mean average path velocity (VAP, μm/s), and other motility parameters. For each sample, on average 100 spermatozoa were analyzed in five fields of view. Sperm concentration was measured using a photometer FEK-M (HV-Lab, RF) instrument. Results of CASA analysis for each of the ejaculates obtained are given in Supplementary Table S1. Each ejaculate sample of fresh semen for all bulls was used for both CASA analysis and IFC analysis for viability assessment with propidium iodide (Sigma-Aldrich, USA) staining. Frozen samples, taken as 0.5 ml straws, were thawed at 37°C, and then analyzed under IFC for both viability and morphological assessments.

### 2.3. Imaging flow cytometry: data acquisition

Since the study was focused on the application of morphological parameters using a deep learning model, the number of images obtained from one ejaculate was sufficient for model training. For each of 6 bulls, there were two fresh semen ejaculates obtained during spring and fall periods and one frozen ejaculate obtained in the summer, resulting in a total of 1.8 million images obtained from IFC. Images were acquired using ImageStream^X^ Mark II (Amnis-Cytek, USA) imaging flow cytometer, equipped with 405, 488, 561, and 642 nm lasers and a brightfield light source, using INSPIRE vs. software (Amnis-Cytek, USA). Images used for neural network training were acquired using a 40x objective. IFC data were initially analyzed using IDEAS vs.6.2 software (Amnis-Cytek, USA). Two files with 50,000 events were recorded for each bull semen sample, and from the initial 1.8 million images, only 436,374 images (n=276,015 for fresh and n=160,359 for frozen) were extracted and further used in manual image labeling and AI algorithm training. The average number of extracted images was about 25,000 for each bull from both groups, as given in Table S2.

### 2.4. Morphology assessment

Obtained images of single-focused spermatozoa were manually classified into the following morphological groups: normal morphology (NM), irregular head shape (IH), abnormal tail (AT), abnormal midpiece (AM), elongated head (EH), proximal cytoplasmic droplets (PCD), distal cytoplasmic droplet (DCD), coiled tail and midpiece (CTM), multiple spermatozoa and debris (16, 37). Representative images of all morphological groups are depicted in Figure 1 below, except for multiple cells and debris groups. The latter two were identified for tidying up the image dataset and removing multiple cells stuck together and debris events, which included fragments of cells and other extracellular components. Only single and focused events with spermatozoa fully fitting into the image frame were utilized for morphological analysis. Images of cells from both fresh and frozen ejaculates were used for model training and classified into morphological groups.

**Figure 1.**
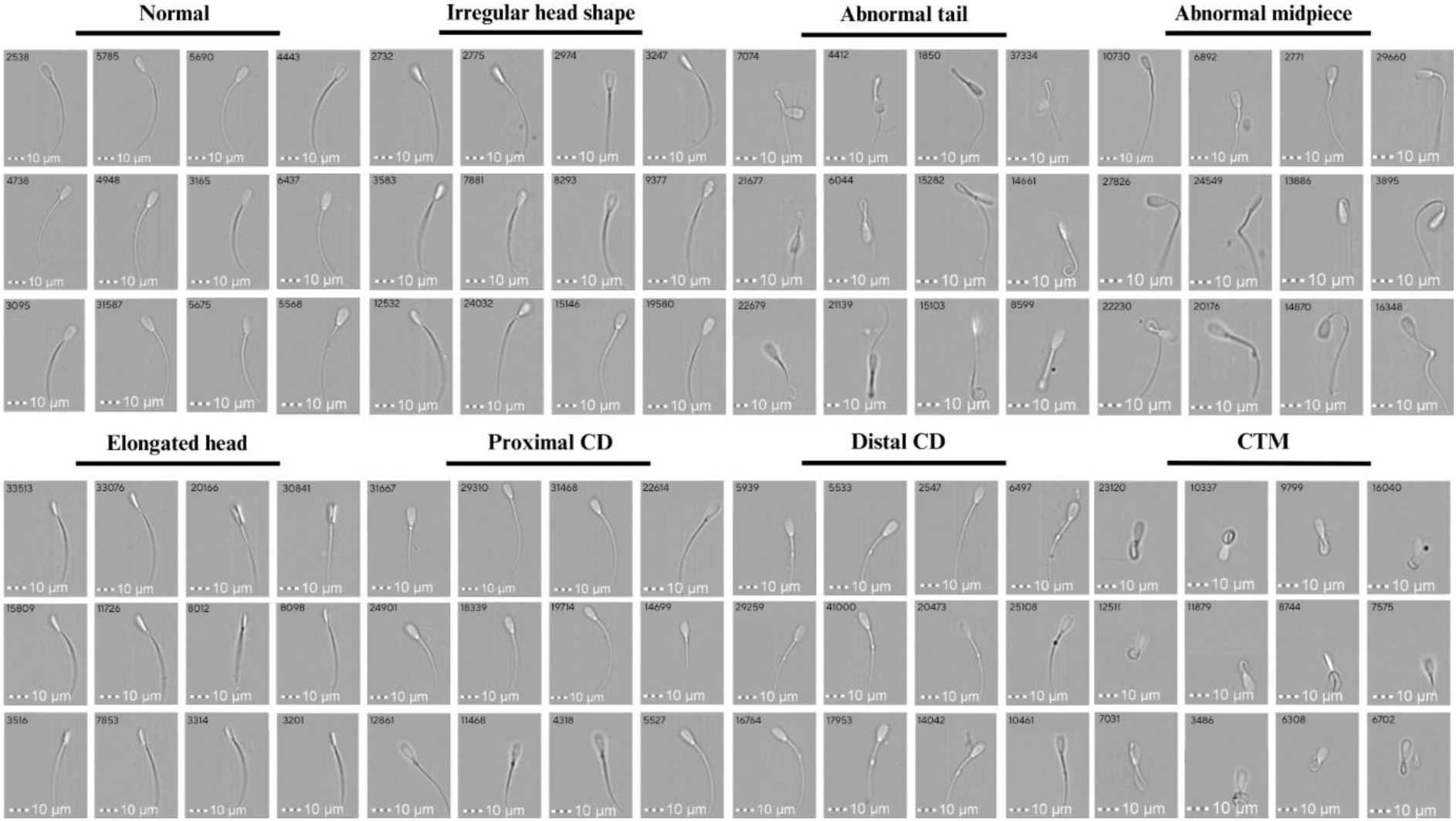
Brightfield images of bovine spermatozoa obtained from Imagestream^X^ Mark II (Amnis-Cytek, USA) (40x magnification). Each of the eight categories was detected and selected for a control set of images manually and divided into the following groups: normal, irregular head shape, abnormal tail, abnormal midpiece, elongated head, proximal cytoplasmic droplet, distal cytoplasmic droplet, and coiled tail and midpiece.

### 2.5. Model training pipeline

Automated spermatozoa image sorting was performed using Convolutional Neural Network (CNN) classifiers. The overall workflow involved dataset preparation, model architecture exploration, training, and evaluation, including specific experiments for cross-condition and cross-breed generalization (Figure 2).

**Figure 2.**
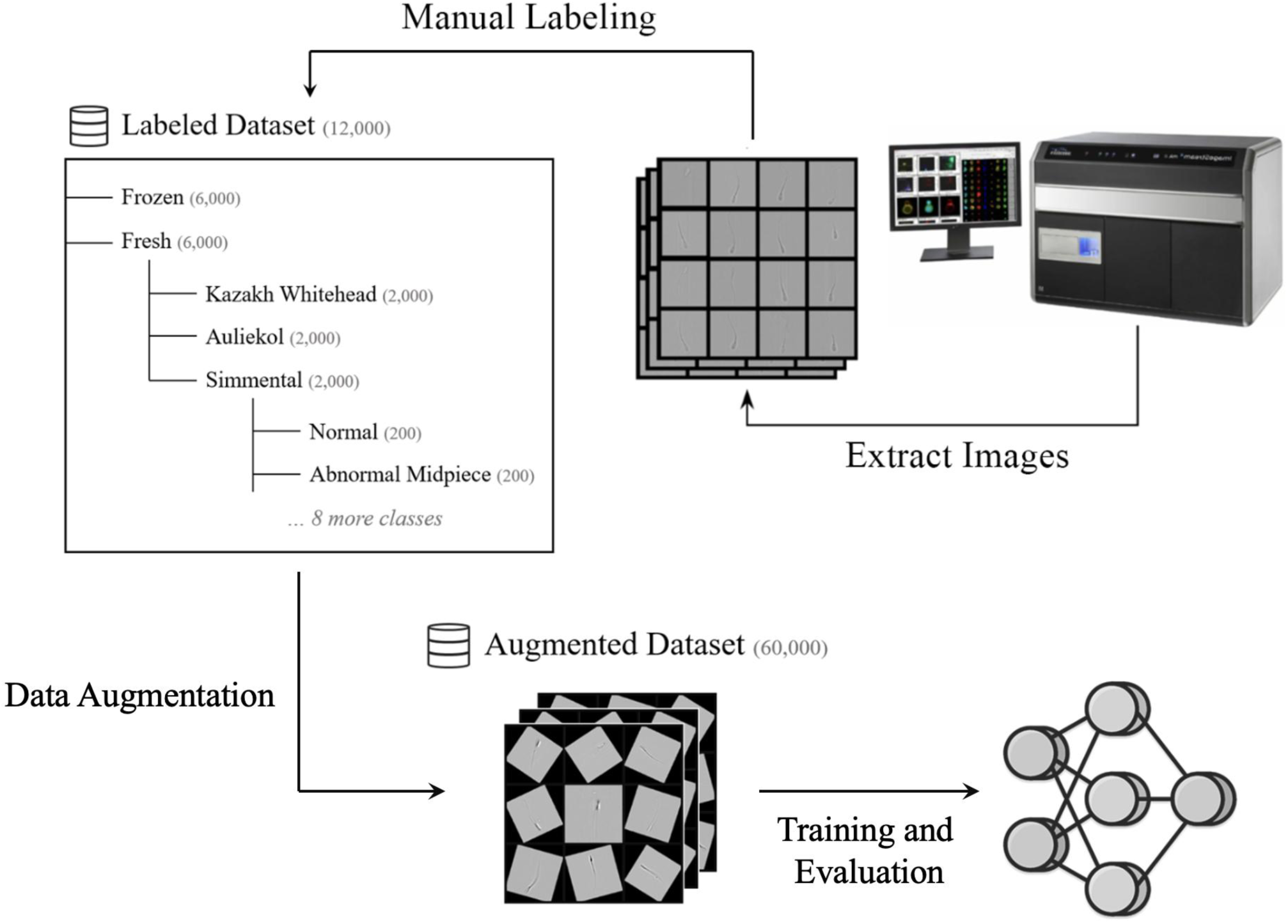
High-level overview of the analysis pipeline for the data selection, processing, and analysis stages. Image collection: the images captured on Imagestream^X^ Mark II (Amnis- Cytek, USA) were manually filtered to constitute focused quality images. Data Augmentation: the images were center-cropped to square ratio and underwent random augmentations, producing 4 additional versions of each image. Training Experiments: multiple models were trained and evaluated in a series of experiments.

#### 2.5.1. Dataset preparation and annotation

Initial data processing involved filtering raw IFC images using IDEAS v.6.2 software (Amnis- Cytek, USA) to obtain images with single-focused spermatozoa. The dataset was iteratively manually filtered to remove debris, partial cells, and significantly blurred images from both fresh and frozen samples across all breeds (Kazakh Whitehead, Simmental, Auliekol).

To construct a balanced dataset suitable for training and robust evaluation, a specific annotation strategy was followed. Before labeling images were standardized through center-cropping and the implementation of a 50×50 pixel size threshold, which removed poorly distinguishable images. High-quality, representative images were manually selected and classified by two independent researchers into ten categories: eight morphological groups (NM, IH, AT, AM, EH, PCD, DCD, CTM) plus categories (’Multiple’ and ’Debris’), following established guidelines (16, 37). The selection aimed to achieve a target of 200 annotated images per category for each of the three bull breeds for both fresh and frozen samples. This annotation resulted in a dataset comprising 12,000 high-quality annotated images (10 morphological classes * 200 images/class * 3 breeds * 2 conditions). As the Kazakh Whitehead constituted four bulls, 50 images per bull were selected for the 10 morphological groups. This dataset exhibits a uniform distribution across classes, breeds, and sample conditions (displayed in Figure 2).

For more reliable model training, four augmented versions were created per original image using random rotations (up to 180°), contrast adjustments (factor 0.3), and random cropping (20%), resulting in a final augmented dataset of 60,000 images used for the experiments described below.

#### 2.5.2. Training strategy

To ensure fair comparison across all experiments, including the generalization tests, a consistent training dataset size was used. For the initial architecture exploration and selection, a subset of 10,000 augmented images was sampled from the primary 60,000-image dataset.

This subset maintained a uniform distribution across conditions, breeds, and classes present in the original 12,000 annotated images. The results are reported in Table 1. Subsequently, for the cross-validation experiments described in Section 2.6, specific datasets of the same size (10,000 augmented images) were curated from relevant subsets of the data (e.g., containing only images from one breed or one condition) while preserving uniform class distribution. This fixed dataset size allows direct comparison of model performance attributable to architecture choices or data source variations. In each experiment, the dataset was divided into 75% / 12.5% / 12.5% proportion for training, validation, and test, respectively.

**Table 1.** Summary of performance results for the tested model architectures and training strategies*.

| Model | Training Strategy | Test Accuracy (%) | Test F1-Score |
| --- | --- | --- | --- |
| YOLOv8 | Standard FT | 90.3% | 0.91 |
|  | LP-FT | 90.6% | 0.89 |
| ResNet-18 | Standard FT | 76.3% | 0.72 |
|  | LP-FT | 83.5% | 0.78 |
| ResNet-50 | Standard FT | 83.2% | 0.78 |
|  | LP-FT | 85.8% | 0.82 |
| MobileNetV3 | Standard FT | 87.9% | 0.87 |
|  | LP-FT | 90.6% | 0.89 |
| **MobileNetV4** | Standard FT | 91.2% | 0.84 |
|  | **LP-FT** | **92.3%** | **0.91** |

\*The best-performed entries are highlighted in bold.

Based on biological imaging application of deep learning models and for diverse experimentation phases, several CNN architectures widely used for classification and segmentation were selected (MobileNetV3, YOLOv8, ResNet-18, ResNet-50, MobileNetV4). They were evaluated using standard fine-tuning (FT) and a two-stage linear probing followed by fine-tuning (LP-FT) protocol (38, 39). The LP-FT strategy involves initially training only the final classification layers while keeping pre-trained ImageNet weights frozen, followed by fine-tuning the entire network with a lower learning rate. All models were trained using the PyTorch framework with GPU acceleration (NVIDIA GeForce RTX 4080), the Adam optimizer, and Cross-Entropy Loss. The training spanned 100 epochs (50 epochs LP, 50 epochs FT) with a batch size of 128, using learning rates of 1e-3 for LP and 1e-5 for FT. Performance was monitored on the validation set after each epoch, and the model checkpoint achieving the highest macro F1-score was saved. The F1-score, the harmonic mean of precision and recall (F1 = 2 * (Precision * Recall) / (Precision + Recall)), balanced the trade-off between false positives and false negatives; the macro average calculates the F1 for each class independently and averages them, treating all classes equally. The best checkpoint for each explored configuration was subsequently evaluated on the held-out test set for a final unbiased performance assessment (Table 1).

## 3. Results

### 3.1. Model training

The comparative performance of different models (MobileNetV3, YOLOv8, ResNet-18, ResNet-50, MobileNetV4) with standard fine-tuning (FT) and the LP-FT strategy was evaluated on the held-out test set (Table 1). The results clearly indicated the superiority of the MobileNet architectures, particularly when using the LP-FT strategy. The ResNet models yielded lower accuracy (below 85%) and showed signs of overfitting. Although YOLOv8 performed well, achieving 90-90.4% accuracy, it was MobileNetV4 paired with LP-FT that achieved the highest overall performance, reaching a test accuracy of 92.3% and a macro F1- score of 0.91. Consequently, MobileNetV4 with LP-FT was selected as the best general- purpose model architecture for this dataset.

The analysis of the confusion matrix for this best model (Figure 3) shows strong classification accuracy across most categories. However, some confusion was noted between classes that have high visual similarity, such as CTM and AT, as well as between EH and IH. There was also minor confusion between the categories ’debris’ and ’multiple cells’. However, this does not impact biological interpretation since both categories are excluded from the analysis. Additionally, the lightweight nature of MobileNetV4 also enabled rapid inference that is suitable for high-throughput analysis.

**Figure 3.**
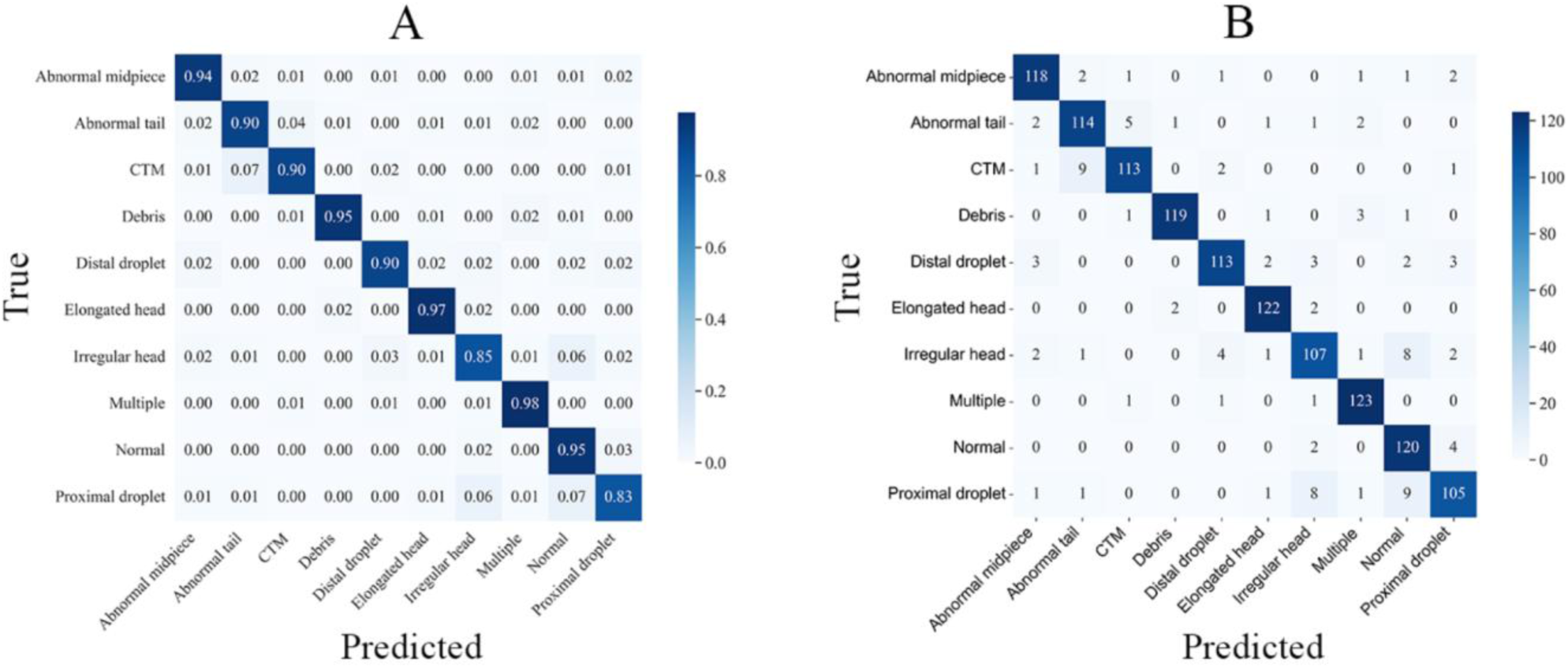
Two confusion matrices illustrate the classification performance of a model across multiple categories. **A.** The normalized confusion matrix displays the same results as proportions (the relative classification accuracy for each category). **B.** The non-normalized confusion matrix shows the absolute counts of predictions for each category, with rows representing the true labels and columns representing the predicted labels. Correct predictions are concentrated along the diagonal.

### 3.2. Cross-validation results

To evaluate the performance transferability across different sample preparations (frozen vs. fresh) and cattle breeds, and to assess potential biases introduced during training, we conducted a series of generalization analyses.

In the first experiment, we trained two models using the selected MobileNetV4 and hyperparameters: one trained exclusively on the aggregated dataset of all frozen samples and another exclusively on the aggregated dataset of all fresh samples. These aggregated datasets were created by combining images from all three cattle breeds for each condition. Each model was then evaluated on images from its own condition (self-evaluation) and from the other condition (cross-condition evaluation).

In the second experiment, the objective was to assess how effectively a model trained on a specific breed within one condition could classify images from other breeds within the same condition. This experiment was conducted separately for the frozen and fresh subsets. Within each condition, three distinct training processes were executed using the chosen architecture and hyperparameters. Each process utilized image data exclusively from one breed: Kazakh Whitehead, Simmental, or Auliekol. Following training, the best checkpoint – determined by the internal validation F1-score - for each single-breed, single-condition model was assessed using images from all three breeds within that same condition. This included both self- evaluation, using the held-out test data from the model’s own breed’s held-out test data, and the cross-breed evaluation, using the held-out test data from the two unseen breeds.

In both experiments, the model performance was measured using overall accuracy, F1-score, detailed per-class metrics (precision, recall, F1-score), and confusion matrices. The results, summarized in Table S3, provide a quantitative assessment of the model’s performance across self-evaluation, cross-condition, and cross-breed scenarios.

#### 3.2.1. Cross-condition performance

Models trained on aggregated data from either frozen or fresh conditions performed well when tested under the same condition. The frozen self-evaluation achieved an accuracy of 88.8% with an F1 score of 0.88 while the fresh self-evaluation reached an accuracy of 90.3% with an F1 score of 0.90. However, when these models were tested under the alternate condition, a performance drop of 6-8% in accuracy was noted. Specifically, the frozen model tested on fresh data achieved an accuracy of 82.2% with an F1 score of 0.82, and the fresh model tested on frozen data attained an accuracy of 82.6% with an F1 score of 0.82. This indicates a domain shift between the sample types in our dataset.

#### 3.2.2. Cross-breed performance

In both frozen and fresh conditions, models that were trained on a single breed consistently performed better when evaluated on the same breed, achieving self-evaluation accuracies between 79% and 87% (Table S3). However, when these models were applied to unseen breeds within the same condition, their performance typically declined, resulting in cross-breed accuracies ranging from 76% to 88.5%. The extent of this decline varied; for instance, the Simmental-trained model showed better generalization, particularly when tested on Kazakh Whitehead, achieving an accuracy of 84.5% (fresh condition). In contrast, the Auliekol-trained model exhibited the most significant challenges, especially in the fresh condition with an accuracy only 75.7% when applied to Kazakh Whitehead. Across all conditions, classifying the IH category consistently proved difficult during cross-breed evaluations. These findings indicate that breed-specific features may impact model predictions, limiting generalization from single-breed training data.

### 3.3. Categorization of sperm images into morphological groups using trained CNN model

Following the completion of CNN model training and cross-validation experiments, the full dataset of 436,374 sperm images (n=276,015 for fresh and n=160,359 for frozen samples) was subjected to classification. After excluding non-suitable images, as outlined in Section 2.5.1, the final number of images that were analyzed using the trained model was 401,535 (n=257,844 for fresh and n=143,691 for frozen). The classification results are summarized in Table S4, with frozen and fresh samples reported separately. Additionally, the fresh semen group was further subdivided by season, encompassing spring and fall.

Head shape anomalies, particularly irregular (IH) and elongated heads (EH) were abundant in both frozen and fresh groups. Morphological abnormalities like distal cytoplasmic droplets (DCD), proximal cytoplasmic droplets (PCD), and abnormal midpiece (AM) were notably present in all of the bulls across all groups. These findings underscore the combined impact of sample preservation method and seasonal variation on sperm quality, which should be carefully considered in reproductive assessments.

## 4. Discussion

The analysis of morphological abnormalities, as well as other semen parameters for assisted reproductive technologies (ART) or in vitro fertilization (IVF), is typically conducted using light microscopes and computer-assisted semen analysis (CASA) systems (40). While most commercial CASA systems can effectively measure sperm motility, they cannot evaluate sperm morphological abnormalities and viability. Several limitations are associated with these techniques. Firstly, they require trained personnel and experts to count and correctly categorize cells into respective morphological groups. Other limitations include a small number of images, low magnification of obtained images, the need for staining to visualize cells, an imbalance between normal and abnormal cell numbers, and increased noise in the resulting images (41).

Spermatozoa morphological abnormalities arise due to several factors, including mechanical damage, oxidative stress, incomplete cell maturation, pathological diseases, and others (42–43), which can affect plasma membrane integrity, viability, motility, mitochondrial membrane potential, DNA composition, and acrosome capacitation (44). Therefore, categorizing these morphological abnormalities is crucial for identifying potential physiological or metabolic changes in the cells and for evaluating the overall fertilizing ability of semen. IFC has proven to be a reliable tool that facilitates robust statistical analysis for the identification of sperm morphology (16, 26, 29–30, 45–46).

Deep learning algorithms allow rapid categorization of cell images into distinct morphological groups (47–48). IDEAS (Amnis-Cytek, USA)-based approach already provides significant opportunities for the analysis of images. Thus, Matamaros-Volante and co-authors (45) used proprietary IDEAS software (Amnis-Cytek, USA) to detect focused images and analyze head and flagella of spermatozoa to study capacitation-induced tyrosine phosphorylation of human spermatozoa. Several deep learning-based approaches were used for sperm quality estimation and classification of abnormal human sperm (49–52). However, only recently suitable databases and AI algorithms were applied for the classification of boar sperm (34, 53). Keller and colleagues (34) applied the IDEAS AI module (Amnis-Cytek, USA) to a deep learning- assisted classification of boar semen into five different morphological groups with 20x, 40x, and 60x magnifications under IFC. For each sample and morphological group, 2,000 images were labeled manually and used for supervised learning for morphological defects and acrosome health status recognition. Based on the study, it was determined that more morphological categories could arise under higher magnification for label-free differentiation of the acrosome status of boar semen.

Thus, Fraczek and co-authors (50) utilized Mask R-CNN for automated segmentation of the sperm head and flagellum from single-channel images. However, their method exhibited low precision when segmenting the spermatozoa tail. Additionally, the size of the database used by some research groups for CNN model’s optimization was limited to 19-20 images, although it included up to 210 single-sperm events (54–55). Hernandez-Herrera and colleagues (56) applied a ResNet50 convolutional neural network to filter out irrelevant images and a second CNN, U-Net 2D, to segment head and midpiece images. However, the above-mentioned research primarily focused on deep-learning algorithms for analyzing human spermatozoa.

In our study, we analyzed a comprehensive dataset comprising 436,374 single-cell, high- resolution images of fresh and frozen spermatozoa from different Kazakhstani bull breeds as well as the widely distributed across Kazakhstan Simmental breed. In a series of experiments, multiple classification strategies and architectures were evaluated using consistent training and data preparation strategies, including MobileNetV3, YOLOv8, ResNet-18, ResNet-50, and MobileNetV4, which was chosen as the final model. The best-performing model, MobileNetV4, was able to achieve an accuracy of 92.3% and an F1-score of 0.91 on a uniformly mixed dataset containing images from various samples and breeds. Utilizing the selected training pipeline, we examined the transferability of learned features across different sample conditions and breeds. As expected, we observed worsening generalization when the data originated from isolated collections such as single breed or bull. Although a fair evaluation of classification methods was attempted, it is worth noting that significantly larger dataset sizes are required to confirm biological bias in domain shift and not personal bias. The cross- validation experiments’ results highlighted a common conclusion: better dataset quality, variety, and size requirements are essential for improved outcomes.

The trained model was used to classify a large dataset of single-cell images (n = 436,374) into 8 morphological groups. On average, approximately 60% of spermatozoa in frozen samples exhibited normal morphology, compared to 34% and 55% in fresh samples collected during the spring and fall seasons, respectively. The proportion of almost all morphologically abnormal sperm cells was reduced. These data are in agreement with findings from other groups regarding the effects of cryopreservation on spermatozoa with abnormal morphology (57). Historically viewed as potentially damaging to sperm, recently, it was suggested that cryopreservation may enhance semen quality by eliminating defective spermatozoa and improving the proportion of healthy spermatozoa, particularly their morphological parameters (58–59). These values are lower than those typically reported in the literature, where the percentage of morphologically normal spermatozoa is expected to be at least 70% when assessed using light microscopy (3, 60). A key distinction of our approach lies in the use of IFC, which enables the high-throughput acquisition and analysis of tens of thousands of individual cell images. In contrast, assessments under light microscopy involve a substantially smaller number of cells, allowing for rapid evaluation but potentially limiting the detection of rare morphological defects, such as tail abnormalities or the presence of proximal cytoplasmic droplets. Furthermore, observer bias is minimized in our method, as morphological classification is performed by a convolutional neural network (CNN) trained on expert-annotated data, ensuring consistent and objective evaluation across all samples.

Advantages of our approach include the analysis across 8 distinct morphological classes and the use of a significantly larger database, the inclusion of leading Central Asian bull breeds alongside the Simmental breed, and the comprehensive analysis of deep learning generalization capabilities of deep learning in the context of spermatozoa cell analysis.

The main limitation of this study was the relatively low number of images in some morphologically abnormal categories. Key areas for improvement include enhancing the size and quality of the database used for CNN training and refinement, as well as the standardization of IFC datasets obtained from different animal breeds. Larger labeled datasets would provide more robust insights into potential biases in cross-validation experiments. Additionally, it is important to note that most modern CNN and vision transformer models can effectively address classification challenges if thorough data preparation and model fine-tuning strategies are performed. We anticipate that advancements in IFC data analysis and a thorough understanding of the cryobiology of spermatozoa will make IFC technology more accessible in the veterinary field and reduce the costs of storage and analysis.

### Author Contributions

A.U. and A.K.: Conceptualization, Data curation, Formal analysis, Investigation, Methodology, Validation; Writing – original draft, Writing – review & editing; B.S.: Conceptualization, Formal analysis, Supervision, Writing – review & editing, Funding acquisition; K.Z., K.T. and M.M.- Formal analysis, Methodology, Writing – review & editing; I.A.V.: Conceptualization, Funding, Supervision, Validation; Writing – review & editing. N.S.B.: Conceptualization, Methodology, Supervision, Project administration; Funding Acquisition; Writing – original draft, Writing – review & editing. All authors have read and agreed to the published version of the manuscript.

## Acknowledgements

We are acknowledging help and advice from Alexander Tikhonov, Nazarbayev University, Vladimir Novokhatsky, Nazarbayev University, and we are grateful to Grigory Marchenko, Amnis-Cytek.

## Institutional Review Board Statement

Not applicable.

## Informed Consent Statement

Not applicable.

## Data Availability Statement

Detailed data are available from the corresponding author by reasonable request.

## Abbreviations

AI -: Artificial Intelligence
AM -: Abnormal Midpiece
ART -: Assisted Reproductive Technologies
AT -: Abnormal Tail
CASA -: Computer-Assisted Sperm Analysis
CNN -: Convolutional Neural Network
CTM -: Coiled Tail and Midpiece
DCD -: Distal Cytoplasmic Droplet EH - Elongated Head
FC -: Flow cytometry
IFC -: Imaging Flow Cytometer
IH -: Irregular Head
IVF -: In Vitro Fertilization
LP-FT -: Linear Probing and Fine-Tuning
NM -: Normal Morphology
PCD -: Proximal Cytoplasmic Droplet
PI -: Propidium Iodide
PM -: Progressive Motility
TM -: Total Motility
VAP -: Average Path Velocity

